# Combined studies of N170/M170 responses to single letters and pseudoletters

**DOI:** 10.1101/2023.10.13.562197

**Authors:** Nima Toussi, Osamu Takai, Sewon Bann, Jacob Rowe, Andrew-John I. Hildebrand, Anthony Herdman

## Abstract

To read efficiently, individuals must be able to rapidly identify letters within their visual networks, which occurs through forming line segments into letters and then letters into words. The temporal processes and utilized brain areas that engage in this process are widely thought to be left-lateralized within the brain. However, a range of studies demonstrate that the processing of unfamiliar stimuli, such as pseudoletters, is temporally delayed and bilaterally processed when compared to letters. The present study investigated the contributions of both hemispheres and how these interactions impact the temporal dynamics of implicit visual processing of single-letters as compared to unfamiliar pseudoletters (false fonts). The results of 5 “in-house” studies are presented within a meta-analysis (synthesis analysis), 3 High-density EEG studies and 2 MEG studies. Of the 3 EEG studies, 2 focused on measuring event related potentials (ERPs) while the participants performed an orthographic discrimination task (letters vs pseudoletters) and the other was a target detection task in which the participants detected infrequent, simple perceptual targets within a series of pseudoletter and letter strings. Of the 2 MEG studies, one was a discrimination task and the other a target detection task. Delayed N170 waveforms to pseudoletters as compared to letters were exhibited across all studies. Lateralization of the ERP differences between letter-evoked and pseudoletter-evoked responses were bilaterally distributed, whereas lateralization measure separately for letters and pseudoletters were primarily left-lateralized. As a whole, these in-house studies indicate that ERPs to letters occur earlier than to pseudoletters, and that interpretation of hemispheric laterality depends on whether the researcher is assessing ERP differences between letters and pseudoletters or the ERP waveforms of the separate letter and pseudoletter conditions.

## INTRODUCTION

Orthographic processing is an indispensable process for reading ability in developing readers(1). Orthographic processing is especially important in developing functional reading proficiency, that is, the ability to identify and understand the meanings of sentences at an efficient rate. Delays in orthographic processing coerce readers into sounding-out words, not only leading to a laborious and choppy reading style, but also diminishing the readers’ comprehension skills(2). Neuroimaging studies investigating orthographic processing using simple letter and word stimuli have provided compelling results regarding the timing and the brain regions involved in this process. Orthographic processing (words, pseudowords, letter strings) appears to begin between 150-170 ms post stimulus presentation; as indicated by event-related potential/field (ERP/ERF) differences among N170/M170 waves evoked by letters (or words), pseudoletters, numbers, and symbols(3–10). This N170/M170 effect is predominantly recorded over the posterior scalp regions. Source-imaging studies of this effect showed involvement of the occipital and inferior-temporal regions(3,5). These results are consistent with the fMRI evidence (11,12). Apart from these general agreements regarding the temporal patterning and anatomical structures involved in orthographic processing, contrasting schools of thought exist regarding the underlying explanations for disparities in EEP/ERF responses and laterality exhibited in orthographic processing.

Prior studies investigating visual processing of word and word-like stimuli using various brain imaging technologies have provided evidence for left-hemisphere lateralization of word-form processing in the primary and secondary visual cortex (13–16). In a broad sense, these studies have found consensus in the finding that visually presented words and pronounceable non-words evoke enhanced activity in the left occipital cortex relative to control stimuli (i.e. pseudoletters and digits). Furthermore, these studies have accumulated evidence for a ‘visual word form’ area, located in the left mid-fusiform gyrus, as the primary area of the brain involved in identifying letters and words from rudimentary shape-like images. These findings imply a predisposition that orthographic processing resides in left-hemispheric brain areas, an idea which is based on the central tenet that language is predominantly processed in the left side of the brain, broadly termed in this paper as the language-dominance hypothesis (17–21).

An improved temporal understanding of these findings have come about from a range of direct and indirect recordings examining the brain’s electrical activity. Intracranial recordings of electrical activity in the inferior temporal lobe have recorded high-amplitude negative responses from the left inferior temporal cortices, termed the N200 for its tendency to peak at 200 ms for orthographic stimuli (words and pseudoletters) and non-orthographic stimuli (faces, objects, etc.). The N200 elicited by all word-like stimuli is greater in the left hemisphere relative to that elicited by faces or objects (22). In extra-cranial EEG recordings, later responses between 200 and 400 ms were found to be greater for pseudoletters than letters (4,10). This apparent processing difference between orthographic and non-orthographic stimuli has been attributed to language-dominant networks in the left inferior temporal cortices used for word reading, in line with the language-dominance hypothesis (10,14,16,23–27).

Similarly, scalp recordings provided by EEG and MEG studies have demonstrated patterns of laterality broadly in-line with intracranial recording reports. The N170, in particular, is an ERP wave that has shown prelexical (word analysis relating to one’s lexicon) sensitivity to orthographic vs. non-orthographic word strings (6,25,28–30). The N170 generally demonstrates greater activity in the left hemisphere for orthographic stimuli but greater activity in the right hemisphere for non-orthographic visual stimuli (31). The left-hemispheric dominance for orthographic stimuli is commonly attributed to letters being a language-based stimulus, which aligns with evidence supporting the hypothesis of a left-hemisphere language dominance.

While a conglomeration of neuroimaging, intracranial recording, and lesion-correlational studies have determined the importance of the left-hemisphere in visual-word recognition, another aspect of orthographic processing may potentially involve the use of a right hemisphere visual-object system, which may support the left-hemisphere Word Form Area by generally processing the perceptual features of visual words, rather than the word stimulus in its entirety (32,33). A growing number of studies are providing contrary findings to larger N170/M170s to letters than pseudoletters and are finding larger and/or more delayed N170/M170s to pseudoletters or symbols than to letters (3–5,34,35). This N170 pseudoletter effect has been localized to generators from bilateral occipital regions (5) or right-hemispheric dominant regions (3,4). Previous work demonstrated that, as compared to letters, pseudoletters or symbols evoked larger N170 amplitudes and, in most cases, more delayed or broader N170 waveforms (3–5,8,35). These findings were likely the cause of difference waveforms between letters and pseudoletters being calculated, whereas many previous studies did not report such difference waveforms.

Furthermore, these findings of larger and delay ERPs to pseudoletters are highly consistent with the literature on visual expertise that showed larger and more delayed N170 amplitudes to unfamiliar (low-level expertise) than familiar (high-level expertise) visual objects such as faces, birds, cars, and learned greebles as well as being consistent with the face-inversion N170 effect (19,36).

The “visual-expertise hypothesis” of orthographic processing revolves around the concept of orthographic processing propagating along the most efficient connections within the brain, rather than being explicitly contained within specialized regions. While the importance of left-hemisphere functions in visual word recognition is difficult to deny, the element of orthographic processing involving a right-lateralized visual-object system which generally processes perceptual features of words rather than their specific orthographic characteristics has led to findings supporting the visual-expertise theory. In addition to the aforementioned evidence originating from the differing results and interpretations arising from N170 studies, behavioral studies using stem-completion word priming have supported the idea of a right-visual object processing area. Marsolek et al., (1992) (37) found that the priming of words based on their visual structure elicited greater effects in the left visual field (and therefore the right hemisphere), indicating that the right hemisphere was more sensitive to discerning the visual specificities of words and letters. Subsequent hemodynamically-based neuroimaging studies (fMRI and PET) have found that while all word-like stimuli activated the lateral extrastriate bilaterally relative to visual fixation tasks, pseudoletters relative to fixation had the greatest degree of right-lateralized extrastriate activation (38,39).

There appears to be inconsistency in the findings among different ERP studies investigating orthographic processing. One way to reconcile inconsistencies is to show results that replicate across multiple studies that use different stimulus sets (e.g., upper vs lower-case stimuli), participant-language groups (e.g., Japanese vs. English), recording methods (e.g., EEG vs. MEG), and study locations (e.g., different lab locations). Observing replicated results among many studies would certainly improve confidence in the findings from any single study. As well, grouping the data across the studies would increase the sample size, thereby improving statistical power.

The goal of this study, therefore, is to address discrepancies which exist in the present literature by combining our multiple studies that investigated ERPs associated with orthographic processing. Although we do not explicitly compare the language-dominance and visual-expertise hypotheses, we do provide consistent evidence across five studies that supports the visual-expertise hypothesis for processing perceptual features of single written characters (i.e., letter or pseudoletter). All studies showed a consistent pattern of results despite being obtained from different electrophysiological measurements (MEG and EEG), different lab environments (multiple sites), different lab personnel collecting the data, different written languages (English and Japanese), different study tasks (categorization and passive viewing), and different stimulus sets (upper- and lower case letters). We anticipate that these results will be applicable in comparing neural processing pathways utilized in reading (i.e. phonological, orthographic, etc.), informing researchers of the interplay between aspects of cognition contributing to reading and the neural mechanisms which produce specific reading behaviors.

## METHODS

### Studies

We included in this “in-house” meta-analysis five adult-participant studies that were conducted across four different labs (MEG lab at The Rotman Research Institute; MEG lab at the Down Syndrome research Foundation, EEG Lab at SFU, and EEG at UBC) using different electrophysiological measurements (EEG and MEG) and recording systems (CTF MEG, NeruoScan EEG, and BIOSEMI EEG) (Table 1). We label this composite study as an “in-house” meta-analysis because we biased the study selection to only those in which the primary author was involved and we had direct access to the data. Thus, this study is not a traditional meta-analysis and holds less credibility than a traditional meta-analysis that has a more inclusive selection of studies. The main reasons for the “in-house” analyses were that we wanted to show consistency across our studies and we had access to all the raw data to allow us to combine the data across the studies in a meaningful way. The names and methodological details for each study are provided below. For the three published studies (4,5,35), we refer the reader to the publications for methodological details and only briefly describe the relevant methods below. Following these study descriptions, we describe how we combined data and performed statistics across the studies (see below, “Combined Analyses” section).

**Table 1.** Brief factsheet on methodologies from each Study utilized in “Combined Analyses”

| Study Number | Recording Method | Sample Size | Task Type | Number of Trials |
| --- | --- | --- | --- | --- |
| Study 1 | EEG | 15 | Mixed<br>(discrimination)<br>task | 288/Letter and<br>Pseudoletter |
| Study 2 | EEG | 11 | Mixed<br>(discrimination)<br>task | 200/Letter and<br>Pseudoletter |
| Study 3 | EEG | 14 | Target task | 200/Letter and<br>Pseudoletter,<br>40/Target |
| Study 4 | MEG | 10 | Mixed<br>(discrimination)<br>task | 288/Letter and<br>Pseudoletter |
| Study 5 | MEG | 10 | Mixed<br>(identification)<br>task | 200/Letter and<br>Pseudoletter |

### Study 1: EEG Active (LetterRhyme)(35)

A main objective of this EEG Active study was to investigate orthographic and phonological processing of single letters. There were three experimental tasks but we only selected results from the orthographic task, “LetterID”, for this paper because it was the most relevant. Please refer to (35) for other results and more details.

#### Participants

Data from 15 English first-language participants (7 females) with normal hearing, vision, and neurological status were included in analyses for the current paper.

#### Stimuli

Letter stimuli were 12 uppercase letters: A, B, D, E, G, H, J, N, P, R, T, and U. Pseudoletter stimuli were created by segmenting and rearranging the line forms of the letter stimuli in order to reduce the differences in the physical properties between letter and pseudoletter stimuli (Fig 1). Letters (288 trials) and pseudoletters (288 trials) were randomly presented for a duration of 500 ms followed by a white fixation dot for a randomized duration of 1250-1750 ms.

**Fig 1.**
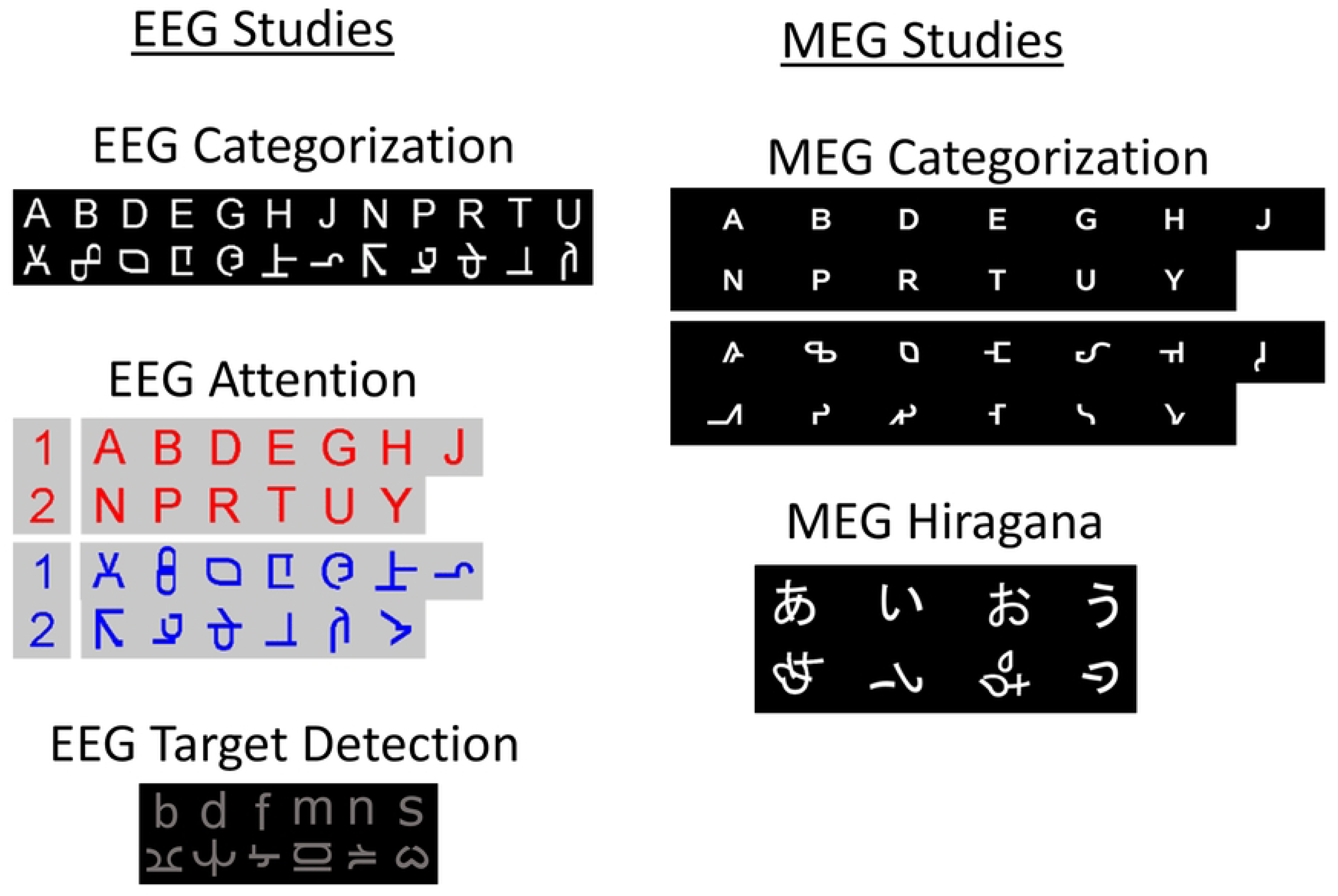
Letter and pseudoletter stimuli for the five studies.

#### Task

Participants were asked to press one button if the stimulus was a letter and a different button if the stimulus was a pseudoletter. We recorded reaction times and accuracies from the participants’ button presses.

#### Recordings

Electroencephalogprahy (EEG) was recorded from participants using an ActiView2 64-channel system (BioSemi, Netherlands) within the BRANE lab at the University of British Columbia. EEG channel data were re-referenced to averaged mastoids. Data were then filtered between 0.1 and 20 Hz and epoched between -500 to 1000 ms relative to stimulus onset. Epoched trials that had amplitudes exceeding ±100 microV between -350 to 850 ms were rejected. Trials were then visually inspected by a trained observer (AH) to ensure that subthreshold ocular artefacts didn’t remain in the data. Those trials that had observable ocular artefacts were also rejected. We feel rejecting all trials that have saccade artefact activity is a more conservative approach as compared to artefact correction. This is because saccades generate activity within visual cortices (40) that would not be removed by using typical PCA or ICA artefact correction procedures. Thus, artefact-related visual activity could still remain within the epoch of interest even after ocular potentials have been corrected by using ICA or PCA correction. We also performed a principal component artifact reduction procedure with a principal component threshold of ±100 microV between -500 to 1000 ms in order to reduce the rising and falling edges of artifacts that might remain within the interval of -350 to 850 ms window (41). This ensured that the artefacts did not contaminate the prestimulus interval during baseline correction between -200 to 0 ms. The artefact-free trials were then averaged together to yield evoked potentials (EPs). We selected the channel that had the greatest N170 response (most negative peak between 120-200ms) over each hemisphere after averaging the ERPs across the group of participants. In this study (EEG Active), the peak channels were PO7 and PO8. We then applied our normalization procedure to these peak channels to obtain the z-transformed responses as defined below in “Combined Analyses”.

### Study 2: EEG Attention Study (LetterTask)(5)

A main objective of this study was to investigate whether paying attention to orthography can affect early EP differences between letters and pseudoletters. Three tasks were used in this study but we selected results from the “Orthography” task for the current paper. Please refer to (5) for more details and other experimental results.

#### Participants

Data from 11 English first-language participants (6 females) with normal hearing, vision, and neurological status were included in analyses for the current paper.

#### Stimuli

13 uppercase letters (A, B, D, E, G, H, J, N, P, R, T, U, and Y) and their pseudoletter counterparts were presented as coloured characters on a light gray background (Fig 1). Letters (200 trials) and pseudoletters (200 trials) were randomly presented for a duration of 500 ms in the central visual field and were followed by a black fixation dot shown for a random duration between 1500 and 2000 ms.

#### Task

Participants were asked to press one of two buttons with their right hand in order to discriminate between letters and pseudoletters. We recorded reaction times and accuracies from the participants’ button presses.

#### Recordings

EEG was collected using a 136-channel BIOSEMI system within SFU’s Human Electrophysiology Lab at Simon Fraser University.

EEG data was filtered between 0.1 and 20 Hz and then epoched between -500 to 1000 ms relative to stimulus onset. Epoched trials that had amplitudes exceeding ±100 microV between - 350 to 850 ms were rejected. Trials were then visually inspected by a trained observer (AH) to ensure that subthreshold ocular artefacts didn’t remain. Those trials that had clear ocular artefacts were subsequently rejected. We also performed a principal component artifact reduction procedure with a principal component threshold of ±100 microV between -500 to 1000 ms in order to reduce the rising and falling edges of artifacts that might remain within the interval of - 350 to 850 ms window (41). This ensured that the artefacts did not contaminate the prestimulus interval during baseline correction between -200 to 0 ms. These artefact free trials were then averaged together to yield the EPs. We selected the channel that had the greatest N170 response (most negative peak between 120-200ms) over each hemisphere after averaging the EPs across the group of participants. In this study (EEG Attention) the peak channels were PO7 and PO8. We then applied our normalization procedure to these peak channels to obtain the z-transformed responses as defined below in “Combined Analyses”.

### Study 3: EEG Target Study (LetterName study)

A main objective of this study was to investigate the brain network dynamics involved audiovisual processing of single letters. Three stimulus conditions (visual only, auditory only, and audiovisual) were used, but we selected results from the “Visual only” condition for the current paper.

#### Participants

Data from 14 adult participants (8 females) with normal hearing, vision, and neurological status were included in analyses for the current paper. Participants were all right handed. English was the primary language for all participants, and all were literate. All participants had normal or corrected to normal (glasses or contacts) vision as tested with a standard Snellen chart.

#### Stimuli

Six lowercase letters (b, d, f, m, n, and s), their pseudoletter counterparts, and a target stimulus (# symbol) were presented as light-gray characters on a black background (Fig 1). The visual stimulus set of letters was limited to six so that their letter-name sounds had similar acoustic properties when being tested in the auditory and audiovisual conditions (data not included in this paper). Letters (200 trials), pseudoletters (200 trials), and targets (40 trials) were randomly presented for a duration of 500 ms in the central visual field and were followed by a light-gray fixation dot shown for a random duration between 1500 and 2000 ms. All stimuli were presented on a 19-inch LCD monitor (DELL/ 1908FPC) set at a distance of approximately 65 cm from the participant’s eyes. A single character covered about 2-3 degrees of vertical and horizontal visual angle.

#### Task

Target Detection. Participants were asked to press one button with their right index finger whenever a target stimulus (#) appeared on the screen. This encouraged participants to pay attention to the stimuli on the screen but they didn’t have to actively discriminate between letters and pseudoletters.

#### Recordings

We performed the exact recordings and analyses as that described in Study #1, EEG Active. In this study (EEG Target Detection) the peak channels found were also PO7 and PO8. We then applied our normalization procedure to these peak channels to obtain the z-transformed responses as defined below.

### Study 4: MEG Active Study (LetterM170)

A main objective of this study was to investigate neural processing of single letters, symbols, and words. There were two experimental blocks (single-characters and strings of characters) with a single task of identifying if the central character is a letter, symbol, or pseudoletter. We only selected results for letters and pseudoletters from the “single-character” block because it is most relevant to this paper.

#### Participants

Data from ten right-handed, adult participants (6 females; ages 19 to 32) with normal hearing, vision, and neurological status were included in this study. English was the primary language for all participants, and all were literate. All participants had normal or corrected to normal (glasses or contacts) vision as tested with a standard Snellen chart. All participants provided informed consent to participate in this study and were paid $10/hour for their participation. Ethics was approved by the Research Ethics Board at Simon Fraser University.

#### Stimuli

Letter stimuli were 13 uppercase letters: A, B, D, E, G, H, J, N, P, R, T, U, and Y. Pseudoletter stimuli were created by segmenting and rearranging the line forms of the letter stimuli in order to reduce the differences in the physical properties between letter and pseudoletter stimuli (Fig 1). Letters (288 trials) and pseudoletters (288 trials) were randomly presented for a duration of 500ms followed white a white fixation dot for a randomized duration of 1250-1750 ms.

#### Task

Participants were asked to press one button if the stimulus was a letter and a different button if the stimulus was a pseudoletter. We recorded reaction times and accuracies from the participants’ button presses.

#### Recordings

Magnetic fields over the whole head were recorded using a 151-channel neuromagnetometer system (CTF-system; VSM-Med Tech, Inc) located at the Down Syndrome Research Foundation in Burnaby, Canada. MEG data were filtered between 0.1 and 20 Hz and then epoched between -500 and 1000 ms. Trials with ocular, myogenic, unknown large (>1.5 pT) artefacts between -350 to 600 ms relative to stimulus onset were manually identified by a trained observer (AH) and rejected. Trials were then averaged together to yield evoked fields (EFs). For each participant’s EF data, we selected one sensor from each hemisphere over the posterior scalp that had maximal EFs peaking within 120 to 190 ms. We classified these as the left and right peak sensors to be used in further analyses for this paper. We then applied our normalization procedure to these peak channels to obtain the z-transformed responses as defined below.

### Study 5: MEG “Hiragana” Study

#### Participants

Data from ten right-handed, Japanese participants (6 females) with normal hearing, vision, and neurological status were included in this study.

#### Stimuli

The visual stimuli were Hiragana graphemes (single letters; あ, ぃ, お, and ぅ) for the phonemes /a/, /i/, /o/, and /u/, and pseudo-graphemes (pseudoletters) made by reconfiguring the letters’ line forms (Fig 1). Letters (200 trials) and pseudoletters (200 trials) were randomly presented for 700 ms with a 300-ms blank black screen followed by a cross-hair at central fixation for a random duration between 2000 to 2500 ms.

#### Task

Participants were asked to wait until the cross hair appeared and then to press one button if the stimulus was a letter and a different button if the stimulus was a pseudoletter. We only measured accuracies and not reaction times because participants were asked to delay their responses.

#### Recordings

Magnetic fields over the whole head were recorded using a 151-channel neuromagnetometer system (CTF-system; VSM-Med Tech, Inc) located at The Rotman Research Institute in Toronto, Canada. MEG data were filtered between 0.1 and 20 Hz (modification from original study) and then epoched between -500 and 1000 ms. Trials with ocular, myogenic, unknown large (>1.5 pT) artefacts between -350 to 600 ms relative to stimulus onset were manually identified by a trained observer (AH) and rejected. Trials were then averaged together to yield evoked fields (EFs). For each participant’s EF data, we selected one sensor from each hemisphere over the posterior scalp that had maximal EFs peaking within 120 to 190 ms. We classified these as the left and right peak sensors to be used in further analyses for this paper. We then applied our normalization procedure to these peak channels to obtain the z-transformed responses as defined below.

### Combined Analyses

#### Sensor Choice

Because this current paper’s objective was to reveal consistencies in findings for difference in the N170/M170 responses to letters and pseudoletters, we selected two channels (one from each hemisphere) to represent the N170 responses interval that have been previously shown to have differences in responses within this interval (4,5,35). To evaluate the consistency of the N170/M170 letter-pseudoletter effect being present across studies, we combined the data from the five studies. To do this, we selected the EEG-electrodes and MEG-sensors that had maximal N170 or M170 responses in each hemisphere across participants in each study.

#### Normalized Evoked Responses (nER)

To combine EPs and EFs across studies, we normalized each participant’s EP (or EF) by dividing the EP (or EF) by the standard deviation of the EP (or EF) amplitudes between 0 to 300 ms for the letter condition for the left-hemisphere peak sensor. We now refer to these responses as normalized evoked responses (nERs). We also calculated difference waves by subtracting nERs for pseudoletters from nERs for letters (letter minus pseudoletters). In addition to traditional averaging of the EPs and EFs as described for the studies above, we performed a N170-locked-EP/EF averaging in order to adjust for task demands, age, and other factors that can shift visual EP/EFs in time (see below for more details). This N170-locked-EP/EF procedure is similar in concept to realigning ERP-locked timing to the button press in order to better resolve motor-evoked potentials.

#### Stimulus-Onset Averaging

For stimulus-onset averaging, we performed the typical ERP/ERF averaging procedure as described above for the individual studies. To combine across studies, we selected artefact-free epochs of -200 to 800 ms that were time-locked to onset of the visual stimuli. These epochs were then averaged and used for further analyses.

#### N170-locked-Response averaging

Because task demands and other factors can shift the visual ERP/ERF peak latencies, we performed another type of averaging that aligns the N170/M170 among studies and among participants. We refer to this procedure as a “N170-locked-Response” averaging which is similar to realigning the ERP’s trial onset times to be at the time of the button presses so that the motor-evoked potentials can be revealed. In our N170-locked response analyses, the ERP onset times were shifted such that the N170 peaks would line up across participants. For our purposes, we shifted each participant’s stimulus-onset averaged data by the difference between 170 ms and a participant’s N170 peak latency for Letters in the left-hemisphere electrode/sensor. We chose the N170, instead of P1 or P2, as a timing anchor because all participants had prominent and easily identified N170/M170 peaks in their evoked responses. For example, if a participant had an N170 peak at 165 ms, then all ERP data for this participant was shifted by 5 ms in the positive time direction. Thus, we aligned all participants’ N170/M170 peak responses to Letters for the left sensor to 170 ms while keeping the relative timing to the ERPs for Pseudoletters and the ERPs in the right sensors the same. The N170-locked ERP method assumes that possible differences between Letters and Pseudoletters are locked to the peak of the N170 for each participant, as we hypothesized. Thus, amplitudes of the difference waves should be larger in the N170-locked ERPs than the traditional-ERP averaging.

One of our hypotheses was that the differences between ERPs/ERFs to letters and pseudoletters might be locked to the responses and not precisely to the stimulus onset. If the main effects are more time-locked to the ERP components than the stimulus, then the difference in nERP/nERFs will be larger for the N170-locked-response averages than the stimulus-onset averages. To test this, we compared each sample of the nERP/nERF difference waves for the stimulus-onset averaging to those for the N170-locked-responses averaging using Student t-tests. This was conducted for left- and right-hemisphere electrodes/sensors. We corrected the t-test results for multiple-comparisons by using the False-Discovery Rate (FDR) correction method (42).

#### Laterality

We also evaluated the laterality of the peak-sensor nERP/nERF by calculating the traditional Laterality Index and a less-traditional Laterality Difference (LD). Both approaches were conducted to allow for a comparison with other studies that have used either the LI or LD measures (7,28). In addition, we calculated the LI and LD across samples between 0 to 300 ms to ascertain which hemisphere had the larger response across this interval. This allowed us to identify time periods when the electrodes/sensors had larger ERP amplitudes in either the left or right hemisphere.

LI was calculated by first calculating the envelope response of the nERP/nERF using a Hilbert transform in order to deal with the bipolar nature of ERPs/ERFs. If we used the raw ERP/ERF amplitude, instead of the envelope response, to calculate the LI (see equation 1 below), then the transition from positive to negative ERP/ERF would result in a flipping of which hemisphere is dominant. For example, if the left hemisphere electrode/sensor had larger P1-N1-P2 amplitudes than the right-hemisphere electrode/sensor, then a positive LI would occur during the positive ERP intervals of P1 and P2 but a negative LI would occur for the N1 interval. Thus, an overlay of ERPs time course or reversing the sign of LI values for negative ERP intervals would be required. In addition, bipolar ERPs should not be used in equation 1 below because the zero-point transition from positive to negative ERPs (or vice versa), where the ERP changes the most, would have very small or zero LI values. To compensate for these caveats, we used the envelope of the ERPs to calculate the LI across samples using the following formula:

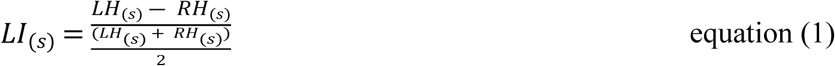

where LH_(s)_ and RH_(s)_ were the amplitudes at sample (s) for the left and right peak sensors, respectively. Thus, positive and negative LI values indicated left- and right-hemisphere laterality respectively.

Because a few previous studies used a left minus right subtraction technique to evaluate hemispheric laterality, we decided to perform a similar analysis for comparison across studies (7,10). We refer to this procedure as the Laterality Difference (LD) but the calculation has one slight alteration with respect to the previously published methods. Because we wanted to observe LD across time samples, we needed a way to deal with the bipolar nature of the ERP. To do this we chose to multiply the difference between LH and RH amplitudes by the sign of the average across LH and RH amplitudes. LD was calculated by using the following equation:

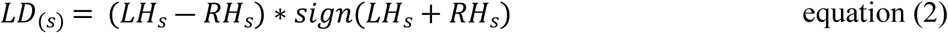

Statistical testing was performed by comparing the LI and LD values for each sample between 0 – 400 ms to a mean of zero using a Student’s t-test. We corrected the t-test results for multiple-comparisons by using the False-Discovery Rate (FDR) correction method using an initial alpha level of .05 (42). Positive LI and LD values reflected significantly larger amplitudes in the left-than right-hemisphere sensor and negative LI or LD values reflected the opposite laterality.

## RESULTS

### N170-locked nERP/nERF Results

The following subsections (P1 interval effects, N170 interval effects, and N170-P2 interval effects) are for organizational and guidance purposes only. We do not imply that the effects found in this study are directly linked to the peaks of the P1, N170, or P2 ERP/ERF components; only that they occur within intervals immediately surrounding these components.

#### P1 interval effects

We found significantly more positive nERP/nERFs surrounding the P1 peak to letters than to pseudoletters (Fig 2 & 3). This was significant only in the right-hemisphere for the grand-study results. The right-hemisphere P1 peaked earlier and larger to letters than pseudoletters in the right hemisphere, which might be causing the significantly more positive nER in the difference wave for this interval. Significant P1 interval effects occurred in the left hemisphere in two of the three EEG studies and none of the MEG studies.

**Fig 2.**
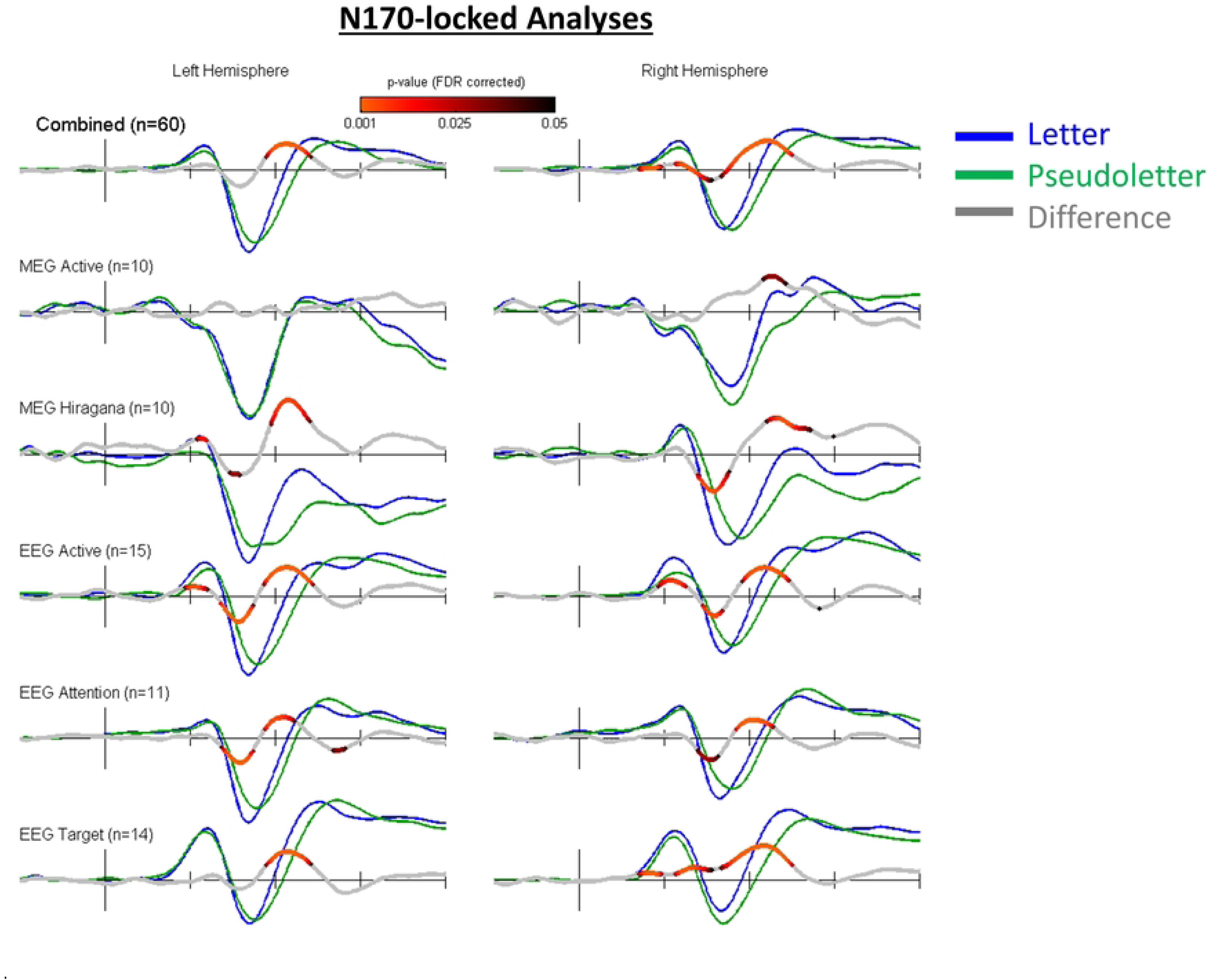
Each Tick on X-axis represents 100ms. Stimulus-Locked waveforms for all five studies. Difference Waveform demonstrates effect calculations between conditions. Combined Analyses (top row of waveforms) shows statistically significant P1 differences for Letters as compared to Pseudoletters and significant differences within the rising and falling limbs of the P1-N170 and N170-P2 responses

**Fig 3.**
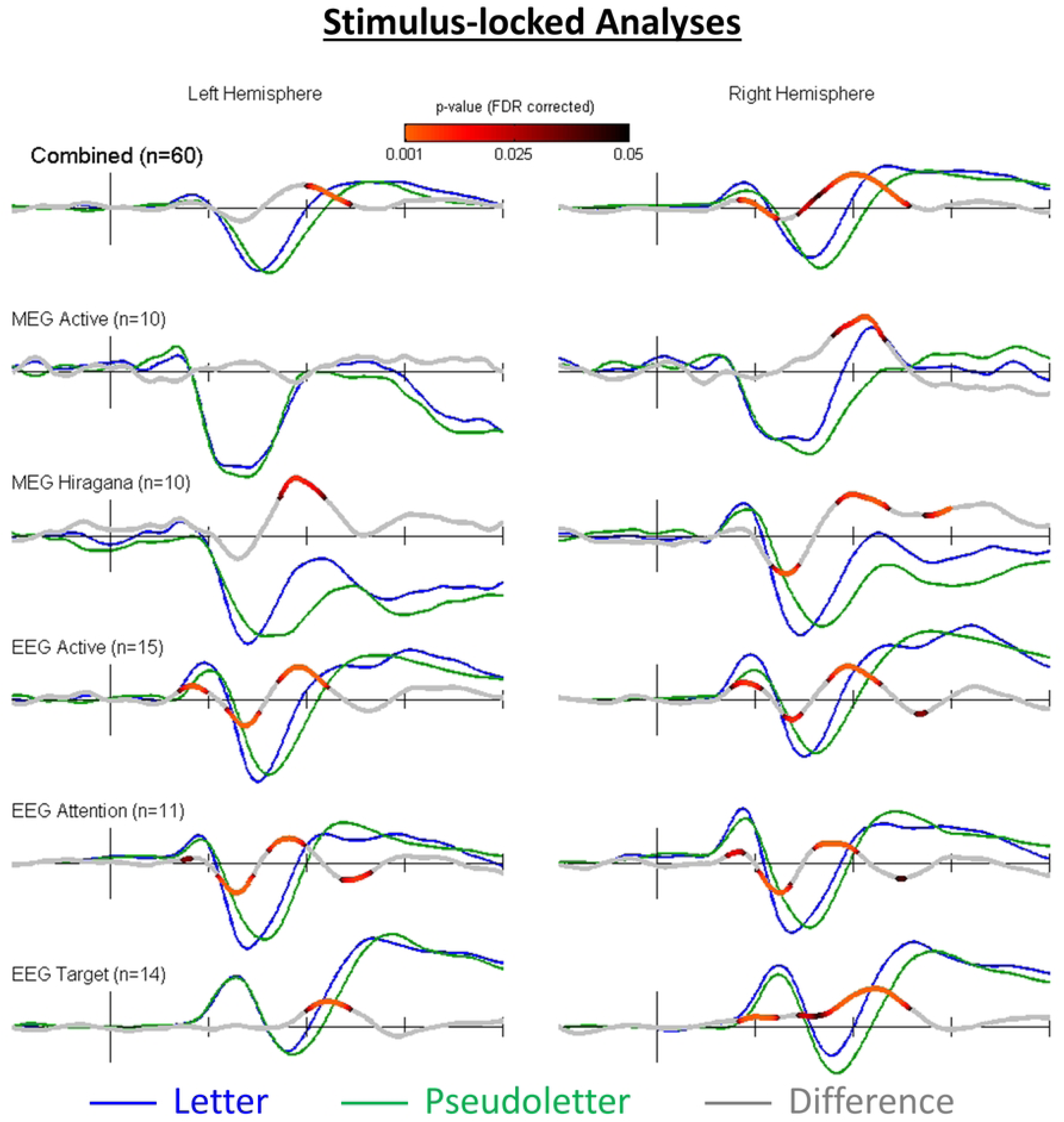
Each Tick on the X-axis represents 100 ms. N170-locked waveforms for all five studies. Difference waveform demonstrates effect calculations between conditions.

These observations are consistent in the non-N170-locked grand-study average of the waveforms with the exception that at the right hemisphere, the letters did not have more negative responses on the falling edge of the N170 than the pseudoletters. In the right hemisphere, the rising phase of the P1 response was significantly larger to letters than pseudoletters.

#### N170 interval effects

The main findings from the meta-analysis of our studies were captured in the grand-study, traditional-averaging waveforms (Fig 3; top panels). The nERPs/nERFs to letters had more negative responses on the falling transition from the P1 to the N170. This resulted in a significantly negative difference wave (letters minus pseudoletter). Although the first P1-to-N170 transition effect is often difficult to see when visually comparing the standard ERPs (blue and green waves), it is clearly apparent (and significant) in the difference ERPs for the left- and right-hemisphere N170-peak electrodes/sensors. This effect becomes even more significant and prominent in the Stimulus-locked analyses (Fig 2).

#### N170-P2 interval Effects

A second N170 effect occurred on the rising edge of the N170. nERPs/nERFs to pseudoletters remained more negative for a longer period during the rising transition from N170 to P2 than did nERPs/nERFs to letters. This led to the largest ERP difference in the difference waves. The peak of this difference did not occur at the peak of the N170, instead it peaked at 185 and 187 ms in the left- and right hemisphere sensors, respectively. This rising-edge N170 effect was more prominent and more significant in the N170-locked than stimulus-locked differences waves (Fig 2 & 3).

Observation of the averaged non-N170-locked waveforms for each study revealed similar trends (more negative responses to letters than pseudoletters on the falling edge of N170, more negative responses to pseudoletters than letters on the rising edge of N170) with a few exceptions. In the LetterName (EEG Target) study, pseudoletters showed a more negative response than letters on the falling edge of the N170, which was statistically significant. A more positive P1 response to letters than pseudoletters was seen across all studies except the MEG Active study (Study 4), where the difference could not be visually observed. Interestingly, this effect was statistically significant only in the LetterTask (EEG Attention) and LetterRhyme (EEG Active) studies. The N170-locked waveforms for individual studies showed very similar statistical results to the non-N170-locked waveforms. Overall, the N170-locked difference waveforms were of larger amplitude than the non-N170-locked difference waveforms.

Observation of the stimulus-locked waveforms within each study revealed similar trends Pseudoletters showed a statistically significant tendency to demonstrate a negative response on the falling edge of the N170 in the LetterName (EEG Target) study. A noteworthy discrepancy between the studies exists regarding the P1 responses; while a more positive P1 response was observed across all studies (except MEG Active study, where the difference couldn’t be visually observed), this effect was statistically significant in only the LetterTask (EEG Attention) and LetterRhyme (EEG Active) studies. Furthermore, there was a statistically significant negative response for letters relative to pseudoletters at P1 in the left hemisphere in the LetterTask and LetterRhyme studies. Although the statistical results for the N170-locked and non-N170-locked waveforms are very similar (within individual studies), the N170-locked waveforms do show greater amplitude than the non-N170-locked waveforms.

### Laterality

The laterality results for grand averaged N170-peak waveform showed that intervals surrounding the P1 and P2 were right-hemisphere dominant, both for letters and pseudoletters (Figs 4 & 5). The same waveforms showed that intervals surrounding the N170 were left-hemisphere dominant for both letters and pseudoletters. Statistically, these effects were significant only for the N170-locked laterality difference waveforms for letters but were observed visually across all grand averaged waveforms (letter, pseudoletter, N170-locked, non-N170-locked). The grand-averaged difference waveforms between letters and pseudoletters showed some right lateralization. In the N170-locked data, this was statistically significant only in the P1 interval, whereas in the non-N170-locked data this was significant in the P1 and N170 intervals. The laterality index data displayed similar results, with the addition of a statistically significant right-lateralized interval in the P2 interval for the letter-pseudoletter N170-locked difference waveform.

**Fig 4.**
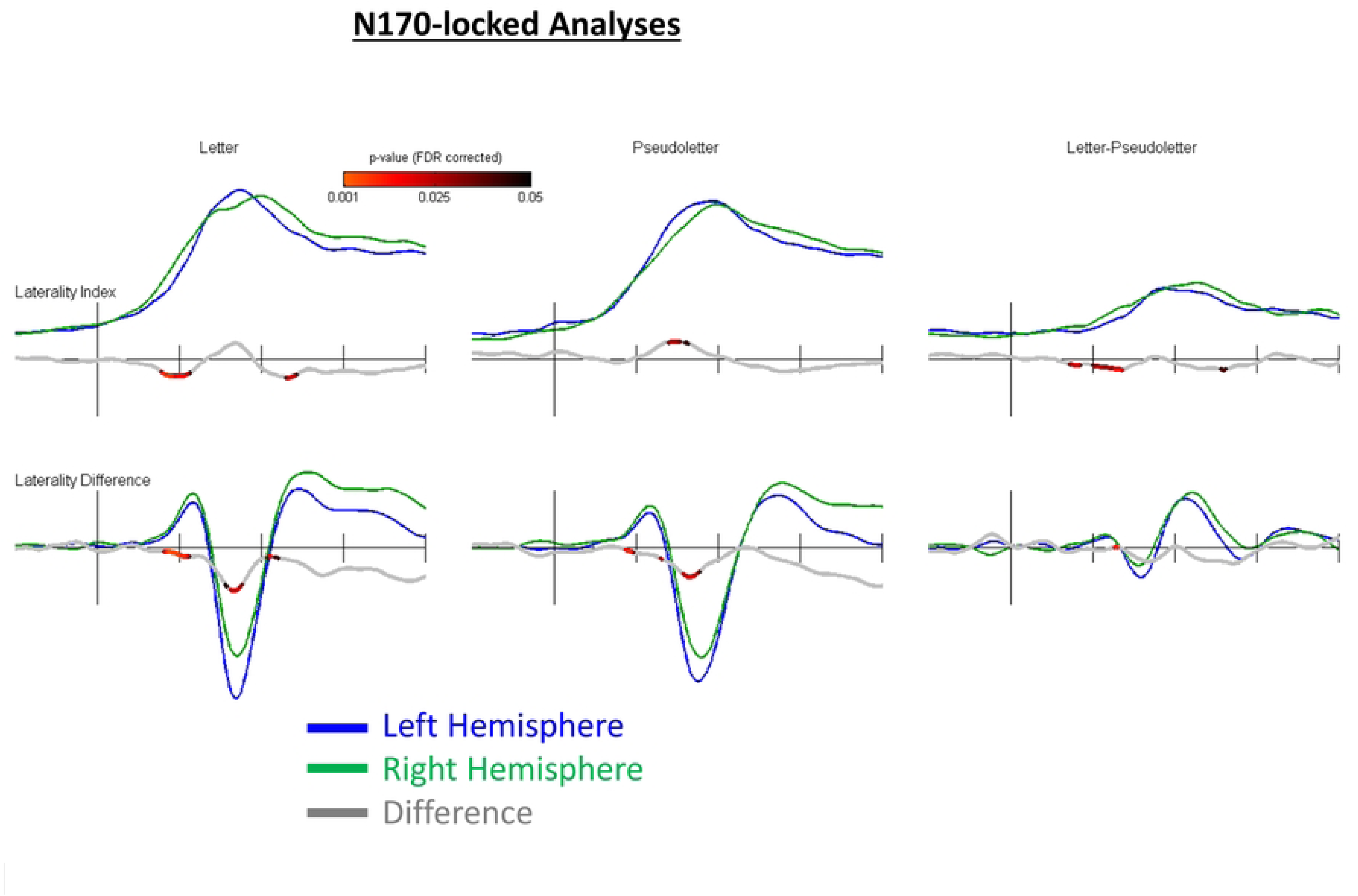
Each Tick on the X-axis indicates 100 ms. Laterality Index and Laterality Difference for N170-locked analyses. Like Fig 2 & 3, grey line indicates difference between Left and Right Hemispheres, with signficant p-values shaded in red/black

**Fig 5.**
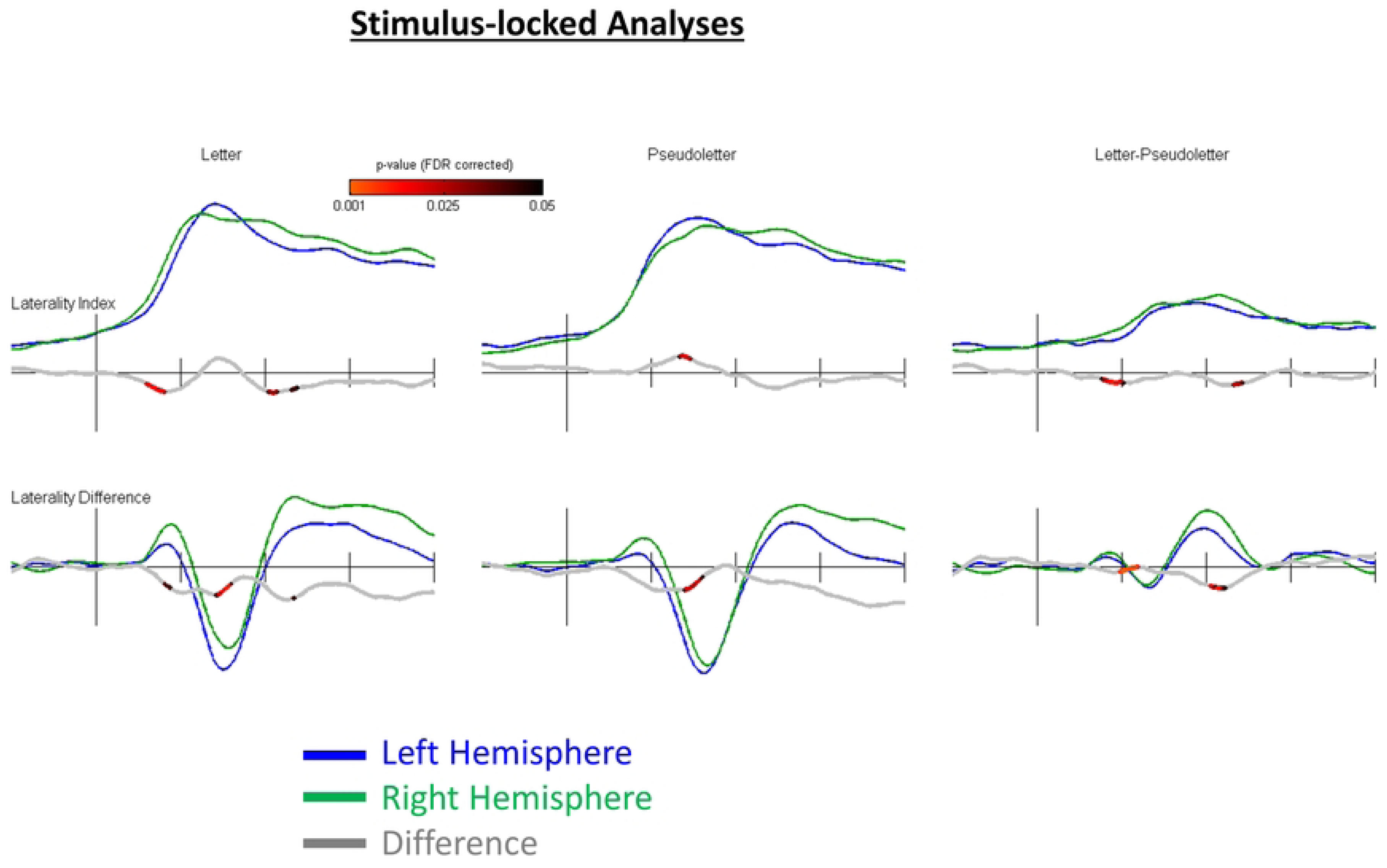
Each Tick on the X-axis indicates 100 ms. Laterality Index and Laterality Difference for Stimulus-locked analyses. General trends similar to those seen in Fig 4, with exception of smaller amplitudes exhibited in Laterality Differences

### Stimulus-onset versus N170-locked-response averaging

The data presented in the LetterName (EEG Target) study showed Stimulus-locked waveforms demonstrated clear similarities to the N-170 locked analysis. Creating the impression that the Stimulus-onset is not event-locked, but rather locked to N170-waveform generation, and hence the stimulus. This challenges past assumptions that it’s a parallel process, time-locked to the N170, rather than impacted by the stimulus.

## DISCUSSION

These studies aimed at investigating the early temporal dynamics and hemispheric laterality of processing single letters and pseudoletters. We found that the N170 to pseudoletters are more delayed and broader as compared to letters. Furthermore, N170s to both letters and pseudoletters were larger in the left than right hemisphere; however, N170 differences between letters and pseudoletters were similar bilaterally with no significant hemispheric dominance.

With reference to the introduction, these findings provide general support for the “visual-expertise” hypothesis. The delayed response to pseudoletters provides support for the idea that the bottom-up processing of the general perceptual features of pseudoletters requires additional neural processing relative to letters because of pseudoletters’ relative lack of familiarity, hence processing times are longer. Bilaterality within N170 responses between letters and pseudoletters provides further support for the “visual-expertise” hypothesis, in so far that it gives credence to the idea that the left hemisphere lacks neural networks specific to processing letters, rather than general perceptual features of letters and pseudoletters. With regards to the scope of these findings within the broader literature on the subject, these findings do generally contrast with the findings of some studies (6,8,10) and provide general support for others (3,4).

The main finding from the combination of these studies, that the ERPs to pseudoletters were more delayed relative to the ERP to letters, was evident for both stimulus- and N170-locked ERP analyses. Difference waveforms made it straightforward to identify the timing of the effects. Our results provide evidence that it is imperative that proper analysis methods are used to fully visualize the effects across the entire interval of interest and not just at the peak responses. These results therefore clarify the timing of pseudoletter ERPs to letter ERPs across the post-stimulus interval and demonstrate that either an additional ERP component is overlaying on the P1-N170-P2 or that a shift in response timing is generating an apparent additional ERP component. This is difficult to determine which is interpretation is correct and further investigation is needed regarding this particular question. For the following discussion we compartmentalized the main ERP differences as P1 effects, N170 effects, and P2 effects, however we only do so out of convenience and readability.

### P1 interval

We found ERPs to be larger for letters than pseudoletters in the P1 interval. This result might indicate differences in basic stimulus parameters such as luminance and spatial frequency between letter and pseudoletter stimuli (43). However, we attempted to control for this effect as best as possible by rearranging the pixels of the letters to generate pseudoletters. Thus, luminance is equated between letters and pseudoletters. This rearrangement does not however equate for spatial frequency and therefore we cannot fully rule out that this might affect the results. There were at least four unique stimulus sets used across these studies which likely would have diluted any effects of differences in spatial frequency between letters and pseudoletters.

An alternative interpretation is that extensive visual experience with letters has enhanced low-level visual networks for earlier processing of familiar objects (i.e., letters) which was reflected in P1 responses as compared to pseudoletters. Evidence has been demonstrated for “early selection” theories of attention with regards to the P1 effect, which stipulate that perceptual inputs are selectively moderated during perceptual processing prior to identification of the stimulus (43). Neuroimaging studies have strongly suggested that initial modulation in ERPs takes place in the posterior extrastriate cortex, at a level where only elementary visual stimulus features are represented (44,45). Should low-level visual networks experience earlier processing for more familiar stimuli, such as letters, it would contribute to the interpretation that it is those low-level networks which contribute to the amplified P1 response in letters.

### N170 interval

Delayed N170/M170 responses to pseudoletters than letters (i.e., pseudoletter effect) were found in all five of our studies. This result could indicate that neural processing for pseudoletters is lagged relative to letter processing. Theoretically, this lag may be attributed to the unfamiliarity of certain stimuli, such as pseudoletters, which results in greater neural analytic processing when compared to more holistic processing of familiar letter stimuli. Some previous studies (6,10) have found peak N170 amplitudes to be significantly different between letters and pseudoletters, though several past papers have established similar findings to the ones observed in this study in so far as that no significant peak N170 amplitude differences were detected between letters and pseudoletters (3,35,46). This result was commonly attributed to low-level visual analysis common to all stimulus letters/strings.

A closer examination of the existing literature on N170 responses presents the possibility that task-types have a significant influence on results. Several studies utilizing N-back task types commonly report observing larger N170 amplitudes to letter than pseudoletter stimuli (10,47,48). N-back tasks typically ask participants to press a button upon seeing ‘N’ number of repetitions of the stimuli. In N-back studies, participants necessarily retain the stimuli in their working memory, which is naturally advantageous to phonological retrieval imposed by letters as compared to the feature-detection processing imposed by pseudoletters. Given that familiar stimuli, like letters, are more established in one’s working memory, these studies broadly posit an expertise effect to account for the greater N170 peaks which are exhibited to letter stimuli as compared to pseudoletter stimuli. Nevertheless, a cohesive theoretical reasoning doesn’t exist to reconcile the results seen in many ‘Target’ task types with the results seen in the selected ‘N-back’ studies. A larger-scale review is likely necessary to develop an improved understanding of the dissonance in results exhibited between the selected ‘N-back’ studies and the results of the five studies in this paper.

### P2 interval

The majority of previous studies show larger P2 amplitudes to pseudoletters than letters (3,4,28,47). Our results also showed that the P2 responses were delayed for pseudoletters than letters. This is likely produced by the delay in the N170-P2 responses which gives an apparent amplitude difference surrounding the P2 responses, at least for our five studies. We do note; however, that previously published studies have shown clearly larger P2 responses to pseudoletters than letters irrespective of whether or not early response delays were evident (3,4,35,47). Interpretations are highly dependent on the methods by which the data are analyzed. Moreover, it is imperative that difference waves are plotted in order to reveal the exact timing of the differences between conditions. Under the language-dominance hypothesis, letters are thought to activate more cortical regions than pseudoletters, which aligns with the results of larger P2 responses. Previous studies have attributed this finding to a lexicality effect, reflecting specialization for word processing, particularly in the visual word-form area (7).

The majority of research conducted with ERPs has used non-stacked average waveforms. Should a research group receive a homogenous group of participants which have more consistent timing in their N170 responses, their results could mirror those seen in N170-locked ERPs, producing greater N1-P2 amplitudes for letters than pseudoletters. This is more likely to occur in labs that predominantly recruit from a homogenous participant body (i.e. psychology students well-versed in doing visual detection tasks). However, if researchers recruit a less homogenous participant group then the results could look more like the non-stacked ERPs (no amplitude effects). While there is a degree of conjecture involved in this explanation for the differences between the N1-P2 amplitudes, it would provide a very possible methodological and parsimonious explanation for past discrepancies in the ERP literature.

### Laterality

A finding that has been reported to support the language-dominant hypothesis is that letters evoke larger N170/M170 over the left than right hemisphere (47). However, other studies have not been able to replicate this finding, or have shown the opposite effect whereby N170/M170 responses differences were larger over the right than left hemisphere (3,4) or were distributed bilaterally (5) (See Table 2). Moreover, few studies have used the traditional measure of laterality, which is to subtract the left and right-side responses then divide by the sum of the left and right responses. Therefore, it can be difficult to fully compare results between different studies. Unfortunately, difference waves are not reported in most prior studies (3,8,31,47) and thus we cannot accurately compare findings among all past studies.

**Table 2.**
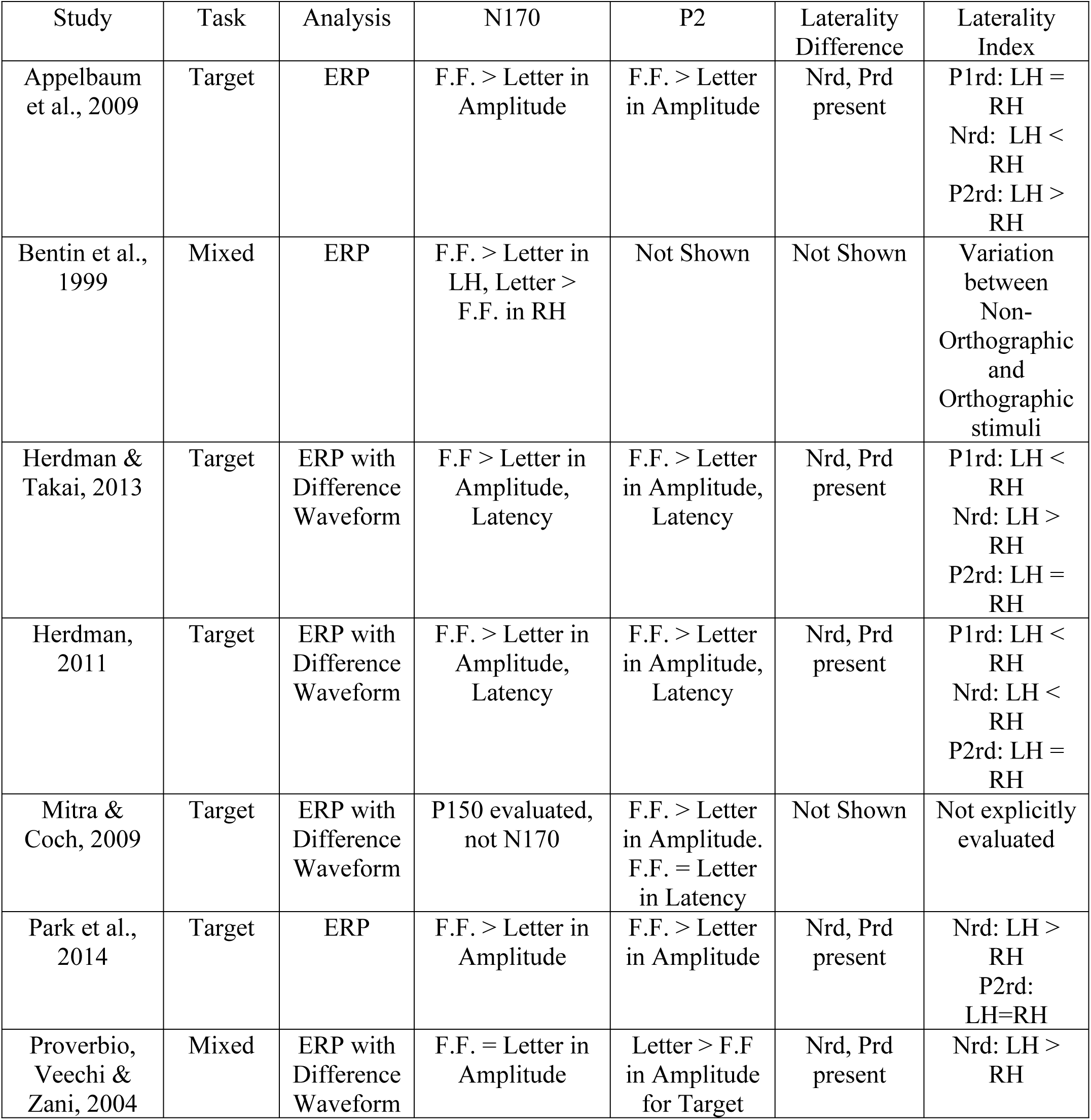
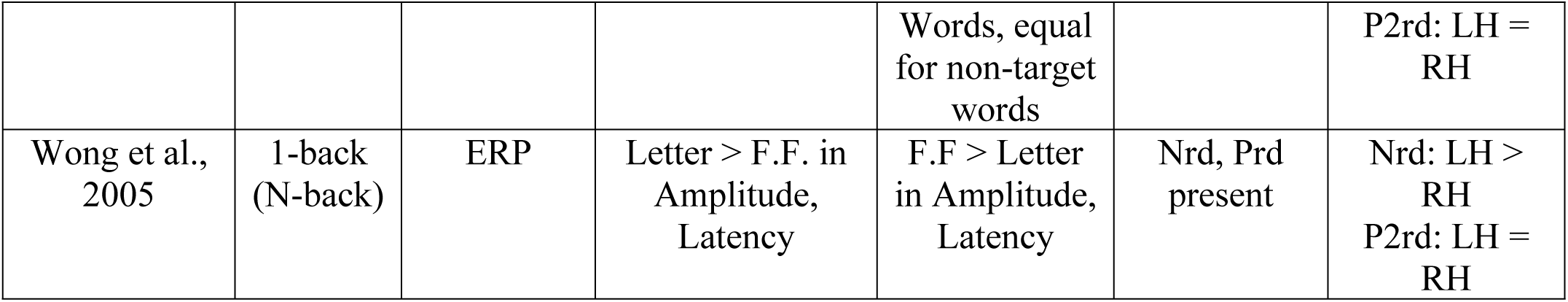
A range of studies presenting varying results on laterality of Negative Response Difference (Nrd) and Positive Response Difference (Prd). Note that studies may refer to N170 and P2 with different terminologies (i.e. RoCC 180 as N170). For brevity, F.F. (False Fonts) is used rather than Pseudoletter within the table.

From both the stimulus-locked and N170-locked results, letter and pseudoletter stimuli evoked larger responses in the left than right hemisphere. This suggests that the left hemisphere is processing the stimuli more in general than the right hemisphere. However, difference waveforms between letters and pseudoletters within the N170 interval showed little hemispheric laterality. This indicates that the both hemispheres are equally involved in the perceptual feature processing of both letters and pseudoletters. Given that the difference waves do not appear to be lateralized, at least for the N170 response, we interpret this as evidence that specialized processing for letters as compared to pseudoletters is bilaterally allocated. Still there were larger responses, in general, in the left than right hemispheric recordings for both letters and pseudoletters. This might indicate a left-dominant processing network involved in orthographic decision tasks.

## CONCLUSION

Evidence was consistent in our “in-house” meta-analysis of two MEG and three EEG studies in that the neural processes involved in discriminating between letters and pseudoletters occurs bilaterally, with an overall left-hemispheric dominance when viewing orthographic (letter) or orthographic-like (pseudoletter) stimuli. The neural delays in processing pseudoletters was also consistent across our five studies and thus we conclude that pseudoletters require more or delayed evaluation of their stimulus features to be able to identify such unfamiliar objects. Overall, a shifting in timing and modulation of ERP amplitudes throughout the post-stimulus interval depends on the stimulus category, letter or pseudoletter.

## ACKNOWLEDGEMENTS

Support of these studies was provided through funding by NSERC, Rotman Research Institute, and the Michael Smith Foundation for Health Research. Special thanks to Dr. John McDonald for use of his lab at Simon Fraser University to undertake Study 2.

## REFERENCES

1. Wagner RK, Torgesen JK, Laughon P, Simmons K, Rashotte CA. Development of young readers’ phonological processing abilities. J Educ Psychol. 1993 Mar;85(1):83–103.

2. Barker TA, Torgesen JK, Wagner RK. The Role of Orthographic Processing Skills on Five Different Reading Tasks. Read Res Q. 1992;27(4):334.

3. Appelbaum L. The temporal dynamics of implicit processing of non-letter, letter, and word-forms in the human visual cortex. Front Hum Neurosci [Internet]. 2009 [cited 2023 Mar 12];3. Available from: http://journal.frontiersin.org/article/10.3389/neuro.09.056.2009/abstract

4. Herdman AT. Functional Communication Within a Perceptual Network Processing Letters and Pseudoletters. J Clin Neurophysiol. 2011 Oct;28(5):441–9.

5. Herdman AT, Takai O. Paying attention to orthography: a visual evoked potential study. Front Hum Neurosci [Internet]. 2013 [cited 2023 Mar 12];7. Available from: http://journal.frontiersin.org/article/10.3389/fnhum.2013.00199/abstract

6. Bentin S, Mouchetant-Rostaing Y, Giard MH, Echallier JF, Pernier J. ERP Manifestations of Processing Printed Words at Different Psycholinguistic Levels: Time Course and Scalp Distribution. J Cogn Neurosci. 1999 May 1;11(3):235–60.

7. Mitra P, Coch D. A masked priming ERP study of letter processing using single letters and false fonts. Cogn Affect Behav Neurosci. 2009 Jun;9(2):216–28.

8. Park J, Chiang C, Brannon EM, Woldorff MG. Experience-dependent Hemispheric Specialization of Letters and Numbers Is Revealed in Early Visual Processing. J Cogn Neurosci. 2014 Oct 1;26(10):2239–49.

9. Proverbio AM, Vecchi L, Zani A. From Orthography to Phonetics: ERP Measures of Grapheme-to-Phoneme Conversion Mechanisms in Reading. J Cogn Neurosci. 2004 Mar 1;16(2):301–17.

10. Wong ACN, Gauthier I, Woroch B, Debuse C, Curran T. An early electrophysiological response associated with expertise in letter perception. Cogn Affect Behav Neurosci. 2005 Sep;5(3):306–18.

11. Gao C, Conte S, Richards JE, Xie W, Hanayik T. The neural sources of N170: Understanding timing of activation in face-selective areas. Psychophysiology. 2019 Jun;56(6):e13336.

12. Yovel G, Sadeh B, Podlipsky I, Hendler T, Zhdanov A. The face-selective ERP component (N170) is correlated with the face-selective areas in the fusiform gyrus (FFA) and the superior temporal sulcus (fSTS) but not the occipital face area (OFA): a simultaneous fMRI-EEG study. J Vis. 2010 Mar 26;8(6):401–401.

13. Cohen L, Dehaene S, Naccache L, Lehéricy S, Dehaene-Lambertz G, Hénaff MA, et al. The visual word form area: Spatial and temporal characterization of an initial stage of reading in normal subjects and posterior split-brain patients. Brain. 2000 Feb;123(2):291–307.

14. Eulitz C, Maess B, Pantev C, Friederici AD, Feige B, Elbert T. Oscillatory neuromagnetic activity induced by language and non-language stimuli. Brain Res Cogn Brain Res. 1996 Sep;4(2):121–32.

15. McCandliss BD, Cohen L, Dehaene S. The visual word form area: expertise for reading in the fusiform gyrus. Trends Cogn Sci. 2003 Jul;7(7):293–9.

16. Miller SL, Wood FB. Electrophysiological indicants of black-white discrimination performance for letter and non-letter patterns. Int J Neurosci. 1995 Jan;80(1–4):299–316.

17. Cai Q, Paulignan Y, Brysbaert M, Ibarrola D, Nazir TA. The Left Ventral Occipito-Temporal Response to Words Depends on Language Lateralization but Not on Visual Familiarity. Cereb Cortex. 2010 May 1;20(5):1153–63.

18. Cohen L. Visual Word Recognition in the Left and Right Hemispheres: Anatomical and Functional Correlates of Peripheral Alexias. Cereb Cortex. 2003 Dec 1;13(12):1313–33.

19. Maurer U, Zevin JD, McCandliss BD. Left-lateralized N170 Effects of Visual Expertise in Reading: Evidence from Japanese Syllabic and Logographic Scripts. J Cogn Neurosci. 2008 Oct 1;20(10):1878–91.

20. Seghier ML, Price CJ. Explaining Left Lateralization for Words in the Ventral Occipitotemporal Cortex. J Neurosci. 2011 Oct 12;31(41):14745–53.

21. Xue G, Dong Q, Chen K, Jin Z, Chen C, Zeng Y, et al. Cerebral asymmetry in children when reading Chinese characters. Cogn Brain Res. 2005 Jul;24(2):206–14.

22. Nobre AC, Allison T, McCarthy G. Word recognition in the human inferior temporal lobe. Nature. 1994 Nov;372(6503):260–3.

23. Cohen L, Dehaene S. Specialization within the ventral stream: the case for the visual word form area. NeuroImage. 2004 May;22(1):466–76.

24. Flowers DL, Jones K, Noble K, VanMeter J, Zeffiro TA, Wood FB, et al. Attention to single letters activates left extrastriate cortex. NeuroImage. 2004 Mar;21(3):829–39.

25. Joseph JE, Cerullo MA, Farley AB, Steinmetz NA, Mier CR. fMRI correlates of cortical specialization and generalization for letter processing. NeuroImage. 2006 Aug;32(2):806–20.

26. Tarkiainen A, Helenius P, Hansen PC, Cornelissen PL, Salmelin R. Dynamics of letter string perception in the human occipitotemporal cortex. Brain. 1999 Nov;122(11):2119–32.

27. Pernet C, Basan S, Doyon B, Cardebat D, Démonet JF, Celsis P. Neural timing of visual implicit categorization. Cogn Brain Res. 2003 Jul;17(2):327–38.

28. James KH, James TW, Jobard G, Wong ACN, Gauthier I. Letter processing in the visual system: Different activation patterns for single letters and strings. Cogn Affect Behav Neurosci. 2005 Dec 1;5(4):452–66.

29. Maurer U, McCandliss BD. The development of visual expertise for words: The contribution of electrophysiology. In: Naples AJ, Grigorenko EL, editors. Single-word reading: behavioral and biological perspectives. New York: Lawrence Erlbaum Associates; 2008. p. 43–63.

30. Tagamets MA, Novick JM, Chalmers ML, Friedman RB. A Parametric Approach to Orthographic Processing in the Brain: An fMRI Study. J Cogn Neurosci. 2000 Mar 1;12(2):281–97.

31. Bentin S, Allison T, Puce A, Perez E, McCarthy G. Electrophysiological Studies of Face Perception in Humans. J Cogn Neurosci. 1996 Nov 1;8(6):551–65.

32. Schendan HE, Ganis G, Kutas M. Neurophysiological evidence for visual perceptual categorization of words and faces within 150 ms. Psychophysiology. 1998 May;35(3):240– 51.

33. Rossion B, Joyce CA, Cottrell GW, Tarr MJ. Early lateralization and orientation tuning for face, word, and object processing in the visual cortex. NeuroImage. 2003 Nov;20(3):1609– 24.

34. Araújo S, Faísca L, Bramão I, Reis A, Petersson KM. Lexical and sublexical orthographic processing: An ERP study with skilled and dyslexic adult readers. Brain Lang. 2015 Feb;141:16–27.

35. Bann SA, Herdman AT. Event Related Potentials Reveal Early Phonological and Orthographic Processing of Single Letters in Letter-Detection and Letter-Rhyme Paradigms. Front Hum Neurosci [Internet]. 2016 Apr 22 [cited 2023 Mar 12];10. Available from: http://journal.frontiersin.org/Article/10.3389/fnhum.2016.00176/abstract

36. Rossion B, Gauthier I. How Does the Brain Process Upright and Inverted Faces? Behav Cogn Neurosci Rev. 2002 Mar;1(1):63–75.

37. Marsolek CJ, Schacter DL, Nicholas CD. Form-specific visual priming for new associations in the right cerebral hemisphere. Mem Cognit. 1996 Sep;24(5):539–56.

38. Petersen SE, Fox PT, Snyder AZ, Raichle ME. Activation of Extrastriate and Frontal Cortical Areas by Visual Words and Word-Like Stimuli. Science. 1990;249(4972):1041–4.

39. Liotti M, Gay CT, Fox PT. Functional imaging and language: evidence from positron emission tomography. J Clin Neurophysiol Off Publ Am Electroencephalogr Soc. 1994 Mar;11(2):175–90.

40. Herdman AT, Ryan JD. Spatio-temporal Brain Dynamics Underlying Saccade Execution, Suppression, and Error-related Feedback. J Cogn Neurosci. 2007 Mar 1;19(3):420–32.

41. Picton TW, Van Roon P, Armilio ML, Berg P, Ille N, Scherg M. The correction of ocular artifacts: a topographic perspective. Clin Neurophysiol. 2000 Jan;111(1):53–65.

42. Benjamini Y, Hochberg Y. Controlling the False Discovery Rate: A Practical and Powerful Approach to Multiple Testing. J R Stat Soc Ser B Methodol. 1995 Jan;57(1):289–300.

43. Hillyard SA, Anllo-Vento L. Event-related brain potentials in the study of visual selective attention. Proc Natl Acad Sci. 1998 Feb 3;95(3):781–7.

44. Desimone R, Duncan J. Neural Mechanisms of Selective Visual Attention. Annu Rev Neurosci. 1995 Mar;18(1):193–222.

45. Ungerleider L. “What” and “where” in the human brain. Curr Opin Neurobiol. 1994;4(2):157–65.

46. Maurer U, Brandeis D, McCandliss BD. Fast, visual specialization for reading in English revealed by the topography of the N170 ERP response. Behav Brain Funct. 2005 Dec;1(1):13.

47. Stevens C, McIlraith A, Rusk N, Niermeyer M, Waller H. Relative laterality of the N170 to single letter stimuli is predicted by a concurrent neural index of implicit processing of letternames. Neuropsychologia. 2013 Mar;51(4):667–74.

48. Xue G, Jiang T, Chen C, Dong Q. Language experience shapes early electrophysiological responses to visual stimuli: the effects of writing system, stimulus length, and presentation duration. NeuroImage. 2008 Feb 15;39(4):2025–37.

